# Structural and Preliminary Biochemical Characterization of MppQ, a PLP-Dependent Aminotransferase from *Streptomyces hygroscopicus*

**DOI:** 10.1101/2022.04.03.486910

**Authors:** Nemanja Vuksanovic, Dante A. Serrano, Alan W. Schwabacher, Nicholas R. Silvaggi

## Abstract

MppQ is an enzyme of unknown function from *Streptomyces hygroscopicus* that is involved in the biosynthesis of the nonproteinogenic amino acid L-enduracididine (L-End). Since L-End is a component of several peptides showing high activity against methicillin-resistant *Staphylococcus aureus* (MRSA), a complete understanding of its biosynthetic pathway is of utmost importance for developing chemoenzymatic routes for syntheses of novel antibiotics. In this work, we report high-resolution X-ray crystal structures of MppQ complexed with pyridoxal-5’-phosphate (PLP) and pyridoxamine-5’-phosphate (PMP). The structure of MppQ shares a fold with known Type I PLP-dependent aminotransferases, consisting of an N-terminal extension, large domain, and a small domain. We also report the first functional characterization of MppQ, which we incubated with enzymatically produced 2-ketoenduracidine and observed conversion to L-End via mass spectroscopy. Additionally, we have observed that MppQ has a relatively high affinity for 2-ketoarginine, a shunt product in the L-End biosynthetic pathway, indicating a possible role of MppQ in increasing efficiency of L-End biosynthesis by converting 2-ketoarginine back to the starting material, L-arginine.

## Introduction

Antibiotic-resistant pathogens are a continuing threat to public health. Mannopeptimycin from *Streptomyces hygroscopicus* and enduracidin from *S. fungicidicus* are non-ribosomally produced peptide antibiotics with activity against a number of Gram-positive pathogens, including drug-resistant strains like methicillin-resistant Staphylococcus aureus (MRSA) and vancomycin-resistant enterococci (VRE) ^1, 2^ Both these agents were discovered decades ago^2^, but despite their promising potency against problem pathogens, they have proved too cytotoxic to be viable therapeutic agents.

The search for less toxic analogues, has been hampered by the difficulty of altering the core peptides of mannopeptimycin and enduracidin since they both contain the unusual non-proteinogenic amino acid L-enduracididine (L-End, **1** in Scheme 1), or its hydroxylated derivative β-hydroxy enduracididine (βhEnd, **2**). This amino acid is not commercially available and, while synthetic routes have been developed^3^, they are multi-step processes starting from an advanced intermediate. The limited availability of this key building block has hampered efforts to make large groups of mannopeptimycin or enduracidin analogs. As a result, we have only a limited understanding of the structure-activity relationships in these compounds. A relatively inexpensive and facile route to L-End would facilitate the search for mannopeptimycin and enduracidin analogs with improved therapeutic properties.

**Scheme 1.**
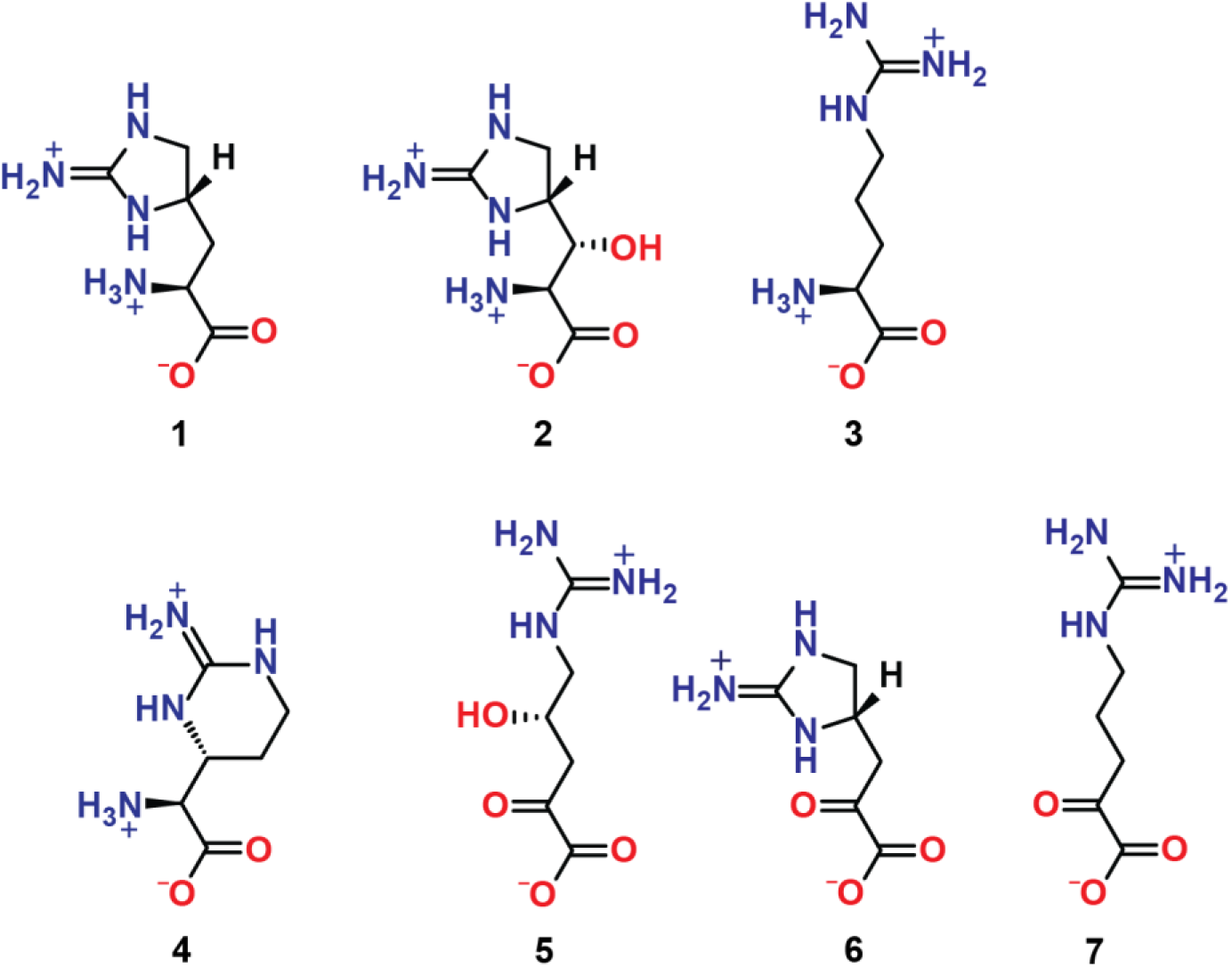
Compounds discussed in this chapter. **1**, L-enduracididine (L-End); **2**, β-hydroxyenduracididine (βhEnd); **3**, L-arginine (L-Arg); **4**, L-capreomycidine (L-Cap); **5**, 2-oxo-4-hydroxy-5-guanidinovaleric acid (i.e. 4-hydroxy-ketoarginine, 4HKA); **6**, 2-ketoenduracididine (2KE); **7**, 2-oxo -5-guanidinovaleric acid (i.e. 2-ketoarginine, 2KA).

Feeding experiments with radio-labelled amino acids^4^ showed that L-End biosynthesis originates with L-Arg (**3**). The biosynthetic clusters for both mannopeptimycin and enduracidin have been characterized^1, 2^, and they share only 3 genes in common: *mpp/endP, Q,* and *R*. Disruption of any of these genes in *S. fungicidicus* abrogates enduracidin production. This defect can be rescued by addition of exogenous L-End to the growth medium.^5^ Thus, the Mpp/EndPQR gene products appear to convert L-Arg to L-End in the biosynthesis of mannopeptimycin and enduracidin, respectively. The mystery of how these three gene products produce L-End was only deepened by comparison to the biosynthetic pathway of a related, L-Arg-derived amino acid, L-capreomycidine (**4**).

Capreomycidine, a component of the antibiotic viomycin, is structurally similar to L-End and is produced by a two-enzyme system. VioC is a non-heme iron- and α-ketoglutarate-dependent oxygenase that hydroxylates L-Arg at the β-carbon atom.^6^ VioD is a fold type I PLP-dependent aminotransferase that catalyzes a unique intramolecular aminotransfer reaction to create the pyrimidinyl ring of **4**.^6^ This system uses one enzyme to activate the β-carbon for cyclization and a second enzyme to create the ring. L-End biosynthesis, by contrast, requires three enzymes. Two of these are PLP-dependent enzymes that could be capable of catalyzing the cyclization step (MppP and MppQ), and none of the three was expected, on the basis of amino acid sequence analysis, to have the catalytic power to activate the γ-carbon atom for cyclization.

The sequence of chemical steps leading from L-Arg to L-End and the intermediates involved became clearer when it was found that MppP from *S. wadayamensis* (SwMppP, a close homolog of the *S. hygroscopicus* enzyme and also in a mannopeptimycin biosynthetic cluster) is a novel type of PLP-dependent oxidase that catalyzes the oxidation of L-Arg to 2-oxo-4-hydroxy-5-guanidinovaleric acid (**5**).^7, 8^ This observation fit in well with previous work on MppR from *S. hygroscopicus* (ShMppR) that indicated this enzyme may operate on an oxidized arginine derivative like **5** to produce the ketone form of L-End (**6**).^9^ The PLP-dependent enzymes are a large group of proteins that catalyze a wide range of reactions, including racemization, decarboxylation, β- and γ-substitution/elimination, and transamination.^10^ This latter reaction accounts for a majority of PLP-dependent enzyme activities. Herein we report the X-ray crystal structure of *S. hygroscopicus* MppQ (ShMppQ), which shows that this enzyme is a fold type I aminotransferase, matching both the tertiary structure and active site architecture of well characterized aminotransferases like aspartate aminotransferase from *E. coli* (EcAAT) and human kynurenine aminotransferase II (1X0M).^11^

The steady state kinetic analysis of the transamination reaction between the possible substrate analog 2-oxo-5-guanidinovaleric acid (**7**) and L-alanine showed that the structural similarities are not surreptitious; ShMppQ catalyzes the aminotransfer reaction. Preliminary kinetic studies involving its physiological substrate, **6**, and L-alanine confirm the role of MppQ as a *bona fide* aminotransferase in the L-End pathway (Scheme 2). Based on the results of mass spectrometry based end-point assays, we have identified L-ornithine as the optimal amino group donor. These studies suggest a likely function for ShMppQ, completing our understanding of L-End biosynthesis.

**Scheme 2.**
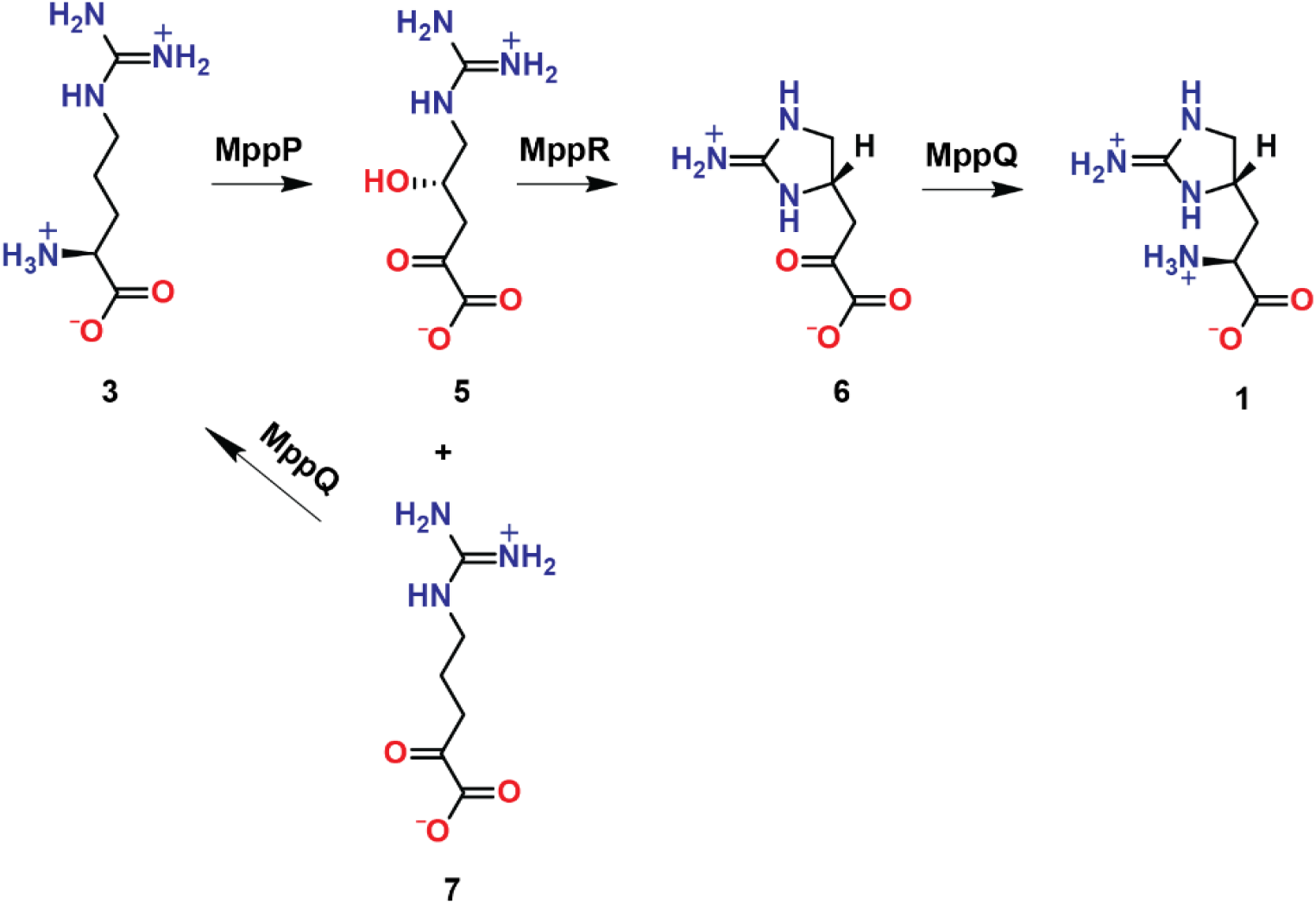
The proposed roles of MppP/R/Q in the biosynthetic pathway of L-End.

## Materials and Methods

### Protein expression and purification

The gene encoding *S. hygroscopicus* MppQ (UniProt accession code Q643B9) was synthesized by GenScript, Inc and subcloned into the pE-SUMOkan vector for expression in *E. coli* BL21 Star cells. Transformed cells were grown at 37 °C in 2-4 L of Lauria-Bertani medium containing 50 μg ml^-1^ kanamycin to an OD_600nm_ of ~1.0. Expression of MppQ was induced with 0.4 mM isopropyl β-D-1-thiogalactopyranoside (IPTG) and the temperature was reduced to 25 °C. After 18 hours, the cells were harvested by centrifugation at 6,000 rpm for 30 minutes. The pellets, approximately 10 g L^-1^ of culture, were resuspended in 5ml of buffer A (25 mM TRIS pH 8.0, 300 mM NaCl, 10 mM imidazole, 200 μM PLP) per gram of cell pellet. The cells were lysed in a circulating ice bath using a Fisher Scientific Model 500 sonic dismembrator at 70% amplitude, for a total of 10 minutes (20 pulses of 30 seconds each, separated by 1-minute rest periods). The resulting lysate was clarified by centrifugation at 39,000 x g (18,000 rpm) for 1 hour. The clarified lysate was passed through a 0.45 μm syringe filter and loaded in 30 ml aliquots onto a 5 ml HisTrap FF column (GE LifeSciences) that had been pre-equilibrated with buffer A (without PLP). After washing with 10 column volumes of buffer A, the protein was washed further and ultimately eluted using a step gradient at 10%, 50%, and 100% buffer B (25 mM TRIS pH 8.0, 300 mM NaCl, 250 mM imidazole). Fractions containing significant amounts of SUMO-MppQ, as judged by a Coomassie-stained SDS-PAGE gel, were pooled and His_6_-Ulp1 protease was added to a final concentration of 2.0 μM. The pooled HisTrap fractions were dialyzed overnight at 4 °C against 3.5 L of buffer C (50 mM TRIS pH 8.0, 150 mM NaCl, 0.5 mM DTT). The following day, the cleaved His6-SUMO tag and protease were removed by passing the dialysate over the HisTrap FF column a second time. The flow-through fractions were collected and judged to be > 95% pure by SDS-PAGE. The protein concentration was quantified by A_280nm_ using the calculated extinction coefficient of 37150 M^-1^ cm^-1^. The protein was dialyzed against 3.5 L of storage buffer (20 mM TRIS pH 8.0, 200 μM PLP) overnight at 4 °C, then concentrated to 11-15 mg ml^-1^ and used directly or snap-frozen in liquid nitrogen.

### Crystallization

Crystals of ShMppQ·PLP (internal aldimine) were grown by the hanging-drop vapor diffusion method over wells containing 15-25 % PEG 3350, and 0.1-0.2 M ammonium citrate trihydrate, trilithium citrate, or ammonium sulfate. Crystals appeared as clusters of thin needles after 3-7 days. The size and morphology of the crystals were improved by microseeding. The crystal used for the MppQ·PMP structure (external aldimine) was grown as above, except that the well solution contained 100mM L-arginine pH 8.0. Crystals were cryo-protected for data collection by soaking briefly in a sequence of solutions containing 30 % PEG 3350, 0.2 M ammonium citrate trihydrate, and 5, 10, or 20 % glycerol.

### Data collection and processing

An initial data set was collected from an unliganded MppQ crystal at 100 K on beamline 21-ID-D of the Advanced Photon Source (Life Science Collaborative Access Team, LS-CAT) equipped with a MAR 300 CCD detector, using an oscillation angle of 1.0 ° over a total of 180 ° with an exposure time of 1 second, a crystal-to-detector distance of 250 mm, and wavelength of 0.97625 Å. Diffraction images were processed using iMOSFLM, POINTLESS, and SCALA as implemented in the CCP4 suite v7.0. This MppQ crystal diffracted to 2.1 Å-resolution. A higher-resolution data set was collected from a different crystal, also at LS-CAT beamline 21-ID-D, using an oscillation angle of 0.5 ° over a total of 140 ° at a distance of 220 mm. The exposure time and wavelength were the same as for the first data set. This crystal was essentially identical to the initial one but was larger and diffracted to 1.8 Å-resolution. These data were also processed using iMOSFLM,^12^ POINTLESS,^13^ and SCALA.^13^ The final data set reported here was collected at LS-CAT beamline 21-ID-G equipped with a MAR 300 CCD detector from an MppQ·L-Arg co-crystal. The X-ray wavelength was fixed at 0.97856, the oscillation range was 0.5 ° over a total of 120 °, the distance was 220 mm, and the exposure time was 1 second. This co-crystal diffracted to 1.5 Å-resolution. These data were processed with the HKL2000 package.^14^ Data processing statistics are given in Table 2-1.

### Structure determination and model refinement

Initial phases were obtained by molecular replacement using PHASER (as implemented in the CCP4 suite v7.0)^15^ with the 2.1 Å-resolution unliganded data set and the structure of *Pyrococcus horikoshii* OT3 kynurenine aminotransferase II (PDB ID 1X0M^11^; 31 % identical) with the 50 N-terminal residues, waters, and PLP-Lysine removed. The lower-resolution data set was used for structure determination, because the “*a*” cell edge of the crystal used to collect the 1.8 Å unliganded data set was oriented parallel to the spindle axis, resulting in poor sampling of the corresponding region of reciprocal space. The Matthews coefficient VM of 2.21 Å^3^ Da^-1^ (41 % solvent) suggested that there were two molecules of MppQ (415 residues, 44.0 kDa) per asymmetric unit. While PHASER was able to find a viable solution (LLG=33, TFZ=9.5), it did not refine well. This poor molecular replacement model was subjected to density modification and automated rebuilding in PHENIX.AutoBuild^16^ to remove bias and improve the model. PHENIX.AutoBuild resulted in a model with 761 residues (740 with side chains placed; ~90 % complete) in 2 chains with R_cryst_ and R_free_ values of 20.0 % and 24.0 %, respectively. This model was used without refinement as the molecular replacement search model to obtain phases for the 1.8 Å-resolution data set (LLG=22,318; TFZ=135). After iterative rounds of restrained refinement in PHENIX.Refine^17^ and manual model building in COOT^18, 19^ the final model of the PLP-bound, internal aldimine form of ShMppQ contained two molecules of MppQ (774 total residues), 816 water molecules, and one molecule of PLP per chain covalently bound at Lys250. The final R_cryst_ and R_free_ values were 14.0 and 17.2 %, respectively.

### Synthesis of 6

The starting material for **6**, 4(S)HKA, was produced enzymatically from L-Arg using the arginine oxidase MppP. A reaction containing 100 mM L-Arg and 200 μM MppP was carried out in 20 mM ammonium acetate, pH 7.0. Bovine catalase (40 mg) was desalted in the reaction buffer (to remove the trehalose stabilizer that would complicate downstream NMR analysis) and added to the reaction mixture to prevent non-enzymatic decarboxylation of 4HKA by the hydrogen peroxide produced by the MppP reaction. The reaction was allowed to proceed overnight at 25 °C while shaking at 150 rpm. The reaction was quenched with 1 volume of methanol and centrifuged at 4,000 rpm for 5 minutes in order to remove the precipitated enzyme. The supernatant was collected and evaporated to dryness at 50 °C in a Centrifan PE Rotary Evaporator. Approximately 1 mg of dried material was resuspended in approximately 1 mL of water, centrifuged at 13 000 rpm for 5 minutes to remove any particulates and analyzed using a ZIC-HILIC column (3.5 μm, 100 Å, 50 x 2.1 mm) paired with Shimadzu LC-MS/MS 8040. Mobile phases used were 20 mM ammonium formate pH 4.0 (designated as “A”) and acetonitrile with 0.1% formic acid (designated as “B”). The analytes were eluted via gradient method at 0.5 mL/min and 35 °C. 15 μL of sample was injected. The following time program was used: hold at 90% B (3.0 min), decrease to 40% B (10 min), hold at 40% B (12 min), increase to 90% B (13 min), hold at 90% B (15). The sample was run in positive ESI mode with a sprayer positioned at 4 mm. The dwell times for products of 190 m/z precursor were set to 80 ms, while dwell time for products of 190 m/z was set to 247 ms. The mass chromatogram (Figure S1) shows the amount of 4HKA and 2KA produced. Note that the peak splitting and distortion has occurred because of column overloading as well as solvent mismatch making the integration less accurate. Based on this data, the sample contains approximately 78% 4(S)HKA and 22% 2KA.

The residue containing a mixture of 4HKA and 2KA (20 mg) was dissolved in a solution containing 180 μM MppR stored in 20 mM sodium phosphate, pH 7.5. The reaction was allowed to proceed for 2 hours at 22 °C on a stir plate set at 100 rpm. The reaction was quenched with 1 volume of methanol and centrifuged at 4,000 rpm for 5 minutes to remove the precipitated enzyme. The supernatant was collected and evaporated to dryness at 50 °C in a Centrifan PE Rotary Evaporator. The reaction products were analyzed using a ZIC-HILIC column (3.5 μm, 100 Å, 50 x 2.1 mm) on a Shimadzu LC-MS/MS 8040. Mobile phases used were 20 mM ammonium formate pH 4.0 (designated as “A”) and acetonitrile with 0.1% formic acid (designated as “B”). The analytes were eluted via gradient method at 0.5 mL/min and 35 C. Injection volume was 0.5 μL. The following time program was used in each run: hold at 90% B (3.0 min), decrease to 40% B (4.5 min), hold at 40% B (6.5 min), increase to 90% B (7.0 min), hold at 90% B (9.0). The sample was run in positive ESI mode with a sprayer positioned at 4 mm. The dwell time for products of 172, 162, and 144 m/z precursors was set to 100 ms. The mass chromatogram (Figure S2) shows that most of 2KE is decarboxylated which most likely occurred during the heating stage at elevated temperature.

### Assay of MppQ activity against 6

Snap-frozen MppQ stored in 50% glycerol, 25 mM TRIS pH 8.5, 200 μM PLP, was thawed and glycerol was removed by buffer exchange with 20 mM TRIS, pH 8.5, 200 μM PLP using a 10 kDa MWCO concentrator. The assay contained a nominal concentration of 50 μM **6** (approximately 10% purity), 40 μM MppQ and 100 mM L-Ala in 20 mM TRIS pH 8.5, 200 μM PLP. The samples were manually quenched at 5, 10, 25, and 50 seconds using 1 volume of 0.1 M HCl. Measurements were conducted in triplicate. The reaction products were analyzed using a ZIC-HILIC column (3.5 μm, 100 Å, 50 x 2.1 mm) on a Shimadzu LC-MS/MS 8040 using the 9 minute gradient described previously. Injection volume was 25 μL for each sample. Dwell time of product ions 172 m/z and 173 m/z was 100 ms.

### Synthesis of 7

The α-keto acid form of arginine was synthesized by enzymatic oxidation of L-arginine using a modified procedure developed by Meister,^20^ and a purification method developed by Stalon et al.^21^ L-Arg (2 g, free base) was dissolved in 50 mL of water and the pH was adjusted using hydrochloric acid to 7.10-7.30. Lyophilized bovine catalase (50 mg) was dissolved in 3 mL of 5 mM sodium phosphate, pH 7.5, followed by buffer exchange using a 10 kDa MWCO concentrator. The sample was concentrated back down to 3 mL. L-Amino acid oxidase (LAAO) from *Crotalus atrox* (60 mg of dried venom) was dissolved in 10 mL water. One half of the catalase solution was added to the LAAO solution. The other half of the catalase solution was added to the L-Arg solution. The LAAO/catalase solution was then combined with the L-Arg/catalase solution, diluted to 100 mL with mqH_2_O and placed in a 1000 mL Erlenmeyer flask and covered loosely with foil. The reaction flask was placed in a shaking incubator at 25 °C, 100 rpm, in the dark for 16 hr. The following day, the reaction was quenched with 100 mL of methanol and centrifuged at 4 °C, 3500 rpm for 10 min. The supernatant was collected. A 30 x 1 cm glass column was packed with 20 g of Dowex 50WX8 200-400 mesh in hydrogen form. A peristaltic pump set to a flow rate of 8 mL/min was used to pass 300 mL of 0.5 M NaOH over the Dowex resin, followed by 300 mL 0.5 M HCl, 300 mL of 0.5 M NaCl, and finally 300 mL of water. The pH of the water issuing from the column must be 7.0 before loading the sample. The supernatant of the LAAO reaction was transferred to the column and the flow-through was collected in a round bottom flask. The column was then washed with 500 mL of water and the flow-through was collected in the same round bottom flask. The flow-through was evaporated to dryness using a rotavap at 40° C. Approximately 1 mg of sample was dissolved in 1 mL of deuterium oxide and centrifuged at 13 000 rpm for 5 minutes. The supernatant was collected and 700 μL of it was loaded into NMR tube. The sample was analyzed by ^1^H, ^13^C and HSQC NMR. In aqueous solution, there are two forms of **7**: linear and cyclic (Appendix A). In the cyclic form the formation of a chiral center at C2 induces diastereotopic properties in the adjacent protons not seen in the linear form. In order to accurately integrate the spectrum, no standard such as acetonitrile was added, as it would overlap with the peaks of interest. The shift alignment was therefore done using the water solvent peak as a reference. This practice is not routinely used, since the shift of exchangeable water protons is dependent on factors such as temperature and pH value.. The solution should not be exposed to pH higher than 8.0 during purification, as this will lead to aldol condensation between two 2-ketoarginine molecules, yielding a conjugated product with a purple color.

### ShMppQ spectrophotometric steady state assay

The steady state kinetics of ShMppQ-catalyzed transamination of L-Ala and **7** was studied by coupling the reaction to *E. coli* lactate dehydrogenase (EcLDH, EC 1.1.1.27). Pyruvate produced in the transamination reaction was reduced by EcLDH resulting in a loss of absorbance at 340 nm due to the oxidation of NADH (ε_340nm_ = 6220 M^-1^cm^-1^). The reactions contained 2 μM ShMppQ, 100 mM L-Ala, 20 U/ml EcLDH, 400 μM NADH, and concentrations of **7** in ranging from 2.5 – 320 μM. The reaction was carried out in 50 mM BICINE pH 7.8, 100 mM NaCl, 50 μM PLP. The absorbance at 340 nm was monitored using an Evolution 300 UV-Vis spectrophotometer (Thermo Scientific). The k_cat_ and KM values were determined from the initial velocity data using Equation 1, where S is the concentration of **7**, v_0_ is the initial velocity, V_M_ is the maximum velocity, and K_M_ is the Michaelis constant. Nonlinear regression analysis was performed in GraphPad Prism.

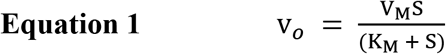

### MppQ Amino Donor Screening

All reactions were carried out in 1 mL volumes of 20 μM ShMppQ, 50 mM bicine, pH 7.8, 150 mM NaCl, and 50 μM PLP. Three identical reactions with 1 mM **3** and 1 mM glyoxylate were manually quenched at 0.5, 1, 2, 5, 10, 15, and 20 minutes by mixing it with 1 volume of pure methanol. Two reactions with 1 mM glycine and 1 mM **7** were quenched in the same manner at 0.5, 10, and 20 minutes. The reactions with glycine and **7** were done in duplicate due to the limited amount of enzyme. The reaction products were analyzed using a ZIC-HILIC column on a Shimadzu LC-MS/MS 8040 using the 9 minute gradient elution method mentioned earlier. Injection volume was 15 μL for each sample. Dwell time for product ion of 174 m/z precursor was 247 ms.

Similar reactions with different amino donor substrates were carried out as end-point assays and contained 10 μM MppQ, 1 mM **7**, 1 mM L-His, L-Gln, L-Orn, L-Ala, L-Phe, L-Ile, or L-Thr, 50 mM bicine, pH 7.8, 10 mM NaCl, and 50 μM PLP. Samples of these reactions were quenched at 5 and 10 minutes with an equal volume of methanol. The reaction products were analyzed using a ZIC-HILIC column (3.5 μm, 100 Å, 50 x 2.1 mm) on a Shimadzu LC-MS/MS 8040 using the 9 minute gradient method mentioned earlier. Dwell time was 100 ms for every product ion resulting from 175 m/z precursor.

## 2.3 Results and discussion

### Overall structure of S. hygroscopicus MppQ

The crystal structure of ShMppQ with the pyridoxal-5’-phosphate (PLP) cofactor covalently bound (internal aldimine) was determined by molecular replacement to a resolution of 1.7 Å (Table 1-1). ShMppQ·PLP crystallized in space group P212121 with unit cell dimensions a = 47.7, b = 114.3, and c = 133.4 Å. The asymmetric unit contains two molecules. There are 14 residues at the N-terminus and 12 from the C-terminus of chain A that are not visible in the electron density. Chain B is similar, with 12 residues at each terminus missing from the model. Chain B also has a break between residues 325 and 329. Superposition of the two chains using Secondary Structure Matching (SSM)^22^ as implemented in COOT, gives a root mean square deviation (RMSD) of 0.32 Å for 382 matched Cα atoms. The two chains in the asymmetric unit form a homodimer (Figure 2-3; 2,974 Å^2^ buried surface area) that matches the quaternary structures observed for known fold type I aminotransferases.^23, 24^ Superposition of the ShMppQ dimer onto the *E. coli* aspartate aminotransferase dimer (PDB ID 1ARS^25^) gives and RMSD of 2.85 Å for 620 of the 775 Cα atoms in the ShMppQ·PLP model.

**Table 1.**
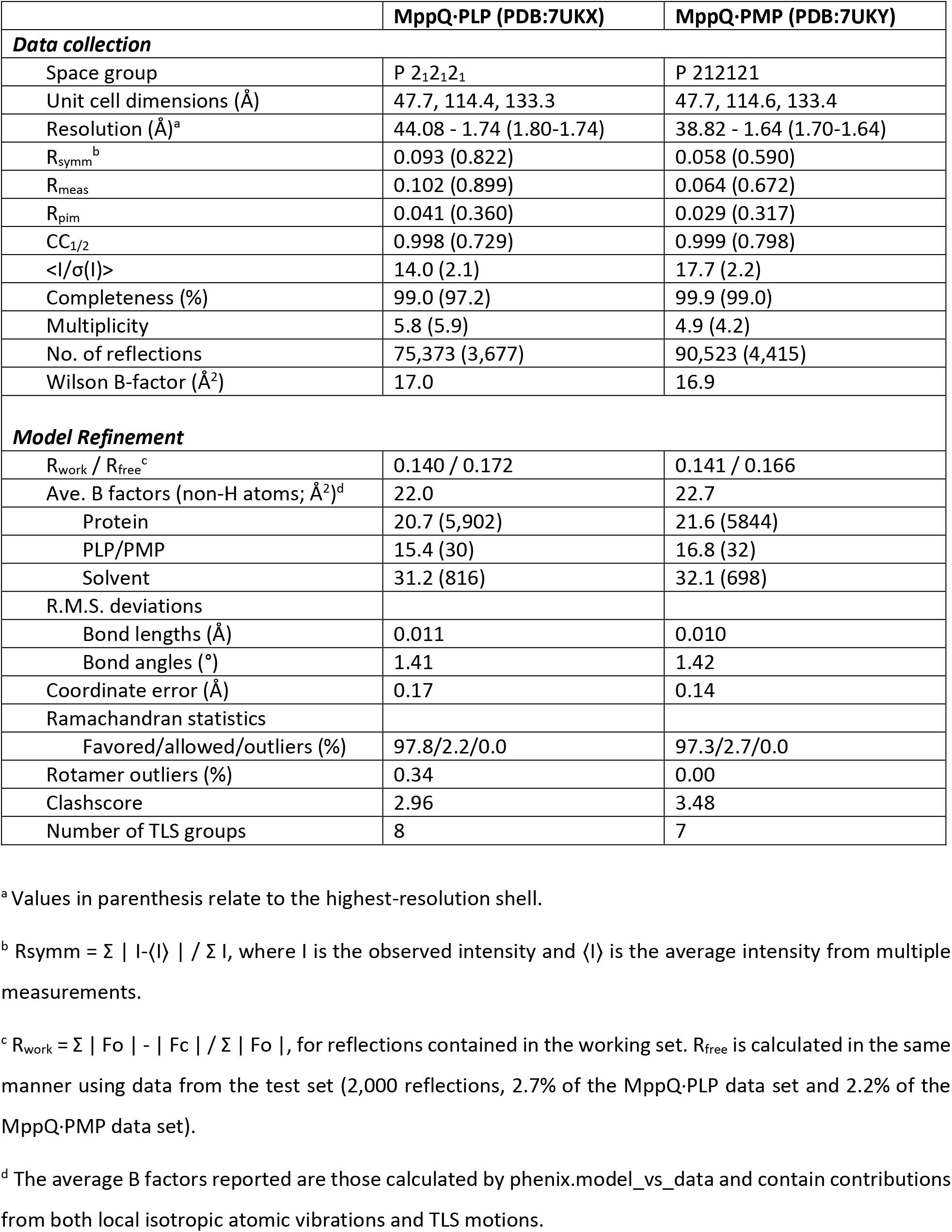
Crystallographic data collection and refinement statistics of MppQ·PLP and MppQ·PMP

The tertiary structure of ShMppQ, like its quaternary structure, matches that of typical fold type I aminotransferases like L-aspartate aminotransferase.^10, 23^ The fold consists of an N-terminal extension (Met1 to Gly35) as well as a large domain (Leu49 to Ala324) and a small domain (Val36 to Leu48 and Met325 to Ser415). The large domain is comprised of a 7-stranded mixed β-sheet flanked by 10 α-helices (α-β-α sandwich), while the small domain has a mixed α-β topology. The large and small domains are linked by a 25-residue-long (~40 Å) helix that serves as the “backbone” of the enzyme. The active site is located at the interface between the large domains of each protomer. As in all known fold type I aminotransferases, the cofactor is held in a cleft between the central β-sheet, and helices α3 and α4 of one subunit, and helix α8 from the other subunit of the dimer (Figure 1).

**Figure 1.**
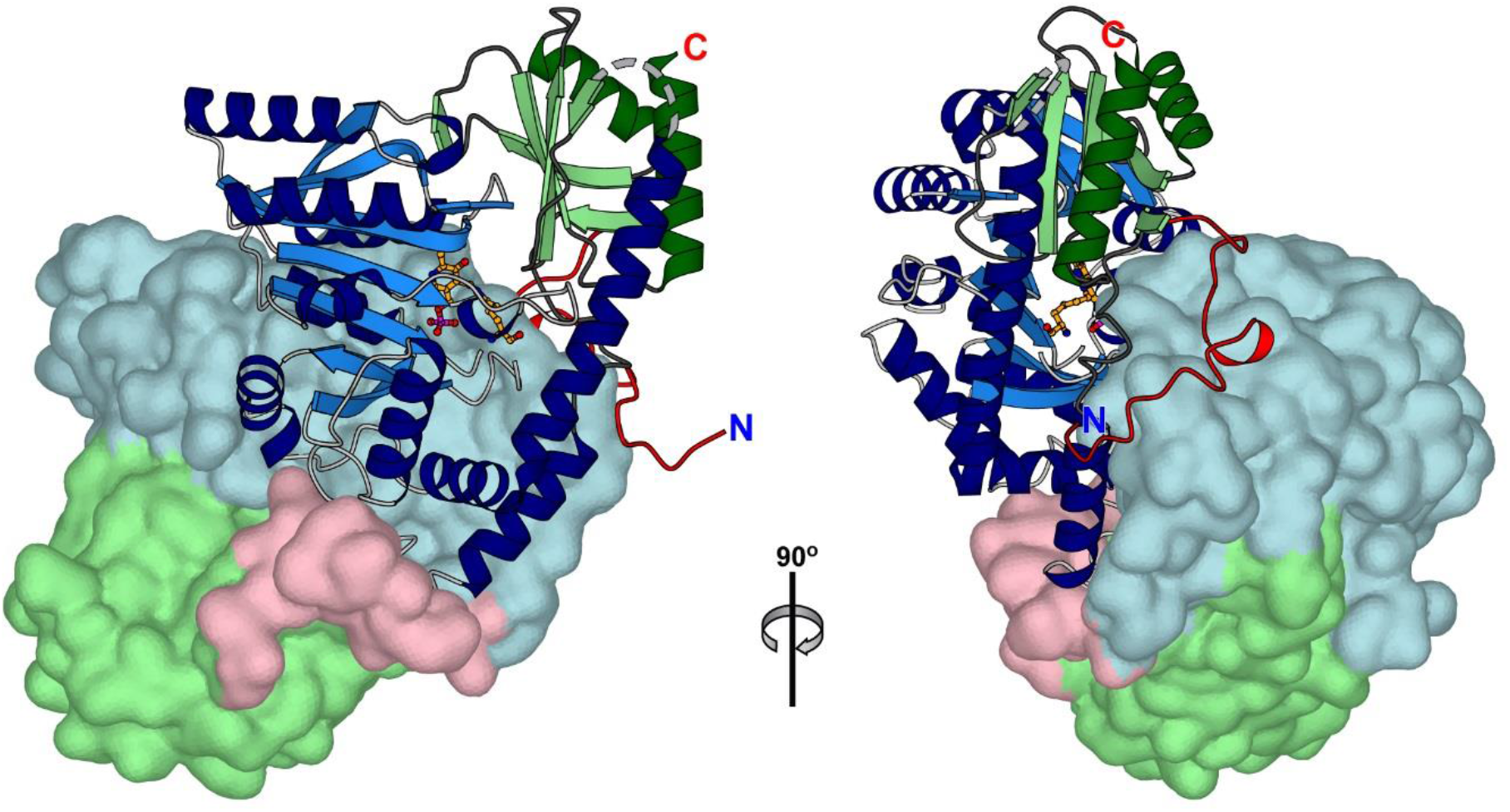
ShMppQ homodimer, with one monomer shown as ribbon representation and other shown as surface representation.

In addition to the *E. coli* aspartate aminotransferase, a PDBeFold^22, 26^ search using chain A of the ShMppQ·PLP model returned 388 unique structures having Q scores of at least 0.14, the threshold value that implies significant structural similarity.^26^ Of these, several had Q scores well above 0.4, implying very strong structural similarity.^26^ These included the human kynurenine aminotransferase II homolog from *Pyrococcus horikoshii* OT3 (PhKAT-II; PDB ID 1X0M^11^; 31 % identical, 2.1 Å RMSD) and the glutamine-phenylpyruvate aminotransferase from *Thermus thermophilus* HB8 (TtGlnAT; PDB ID 1V2D^27^; 29 % identical, 2.2 Å RMSD). The former catalyzes the production of kynurenic acid and glutamate from kynurenine and α-ketoglutarate,^28^ while the latter has been shown to catalyze aminotransfer between a number of aromatic amino acids and α-ketoglutarate.^27^

### Comparison of PLP- and PMP-bound MppQ

The Structure of ShMppQ·PLP shows clear electron density for the covalent attachment of PLP to the catalytic Lys250. The side chains surrounding the cofactor are typical of aspartate aminotransferase (Figure 2). The residues N187 and Y218 are hydrogen bonded to the oxygen atom attached to the pyridinium ring of PLP. D215 forms a salt bridge with the nitrogen of the pyridinium, thus stabilizing it and allowing PLP to act as an electron sink in the reaction with the substrate. The phosphate group of the PLP is within hydrogen bonding distances of S109, R257, S249, S247, as well as Y72 of the other monomer chain (Figure 2C). Most side chains lining the active site in EcAAT active site are identical (Y225, N194, D222, R266, S257, S255), except for subtle differences (PDB Entry: 1ARS).^25^ In EcAAT, it is Trp140 which forms interactions with the PLP aromatic ring via pi-pi stacking, while in MppQ, Y133 if found in this location. EcAAT also has T109 interacting with the phosphate moiety instead of S109.

**Figure 2.**
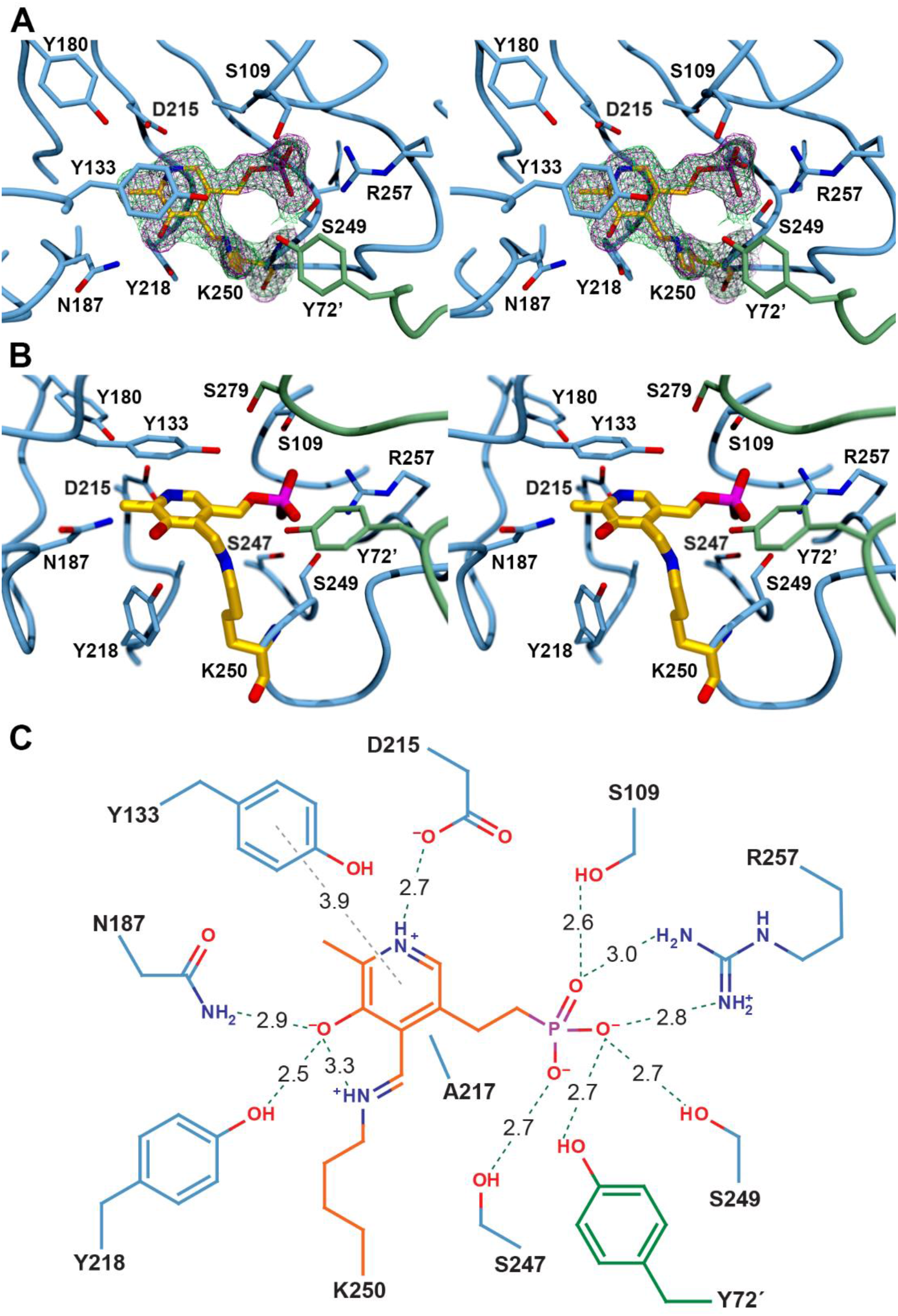
The 2|F_o_|-|F_c_| (magenta mesh) and simulated annealing composite omit (green mesh) electron density maps are shown for the K250-PLP adduct at 1.2 σ (map radius 2.5 Å) in a stereoview of the ShMppQ active site (A). A second stereoview of the active site (B) is shown without electron density and in a different orientation (rotated ~60 ° in the horizontal) to give a clearer picture of the linkage between K250 and the cofactor. A schematic representation of the active site (C) shows the interactions of active site functional groups with the cofactor and each other (associated distances are given in Å).

Co-crystallization of the PLP-bound enzyme with 10 mM L-Arg resulted in trapping of the noncovalent complex between ShMppQ and pyradoxamine-5’-phosphate (PMP). The PMP-bound form of the enzyme was isomorphous with the ShMppQ·PLP complex. These crystals diffracted to 1.6 Å resolution and contained two molecules in the asymmetric unit, arranged in the same quaternary structure observed for the PLP-bound form of the enzyme (0.295 Å RMSD). The disordered regions in each chain are identical to those seen in the ShMppQ·PLP model.

The active site clearly contains PMP, as evidenced by the lack of continuous electron density to Lys250 (Figure 3). Aside from the presence of PMP in the active site, there is virtually no difference between the active sites of the PLP- and PMP-bound forms of ShMppQ. It will be interesting to find out if the change to a “closed” conformation (rotation of the small domain relative to the large domain) observed in some ATases does not happen in MppQ, or if the crystal simply captured an “open” conformation in each case.

**Figure 3.**
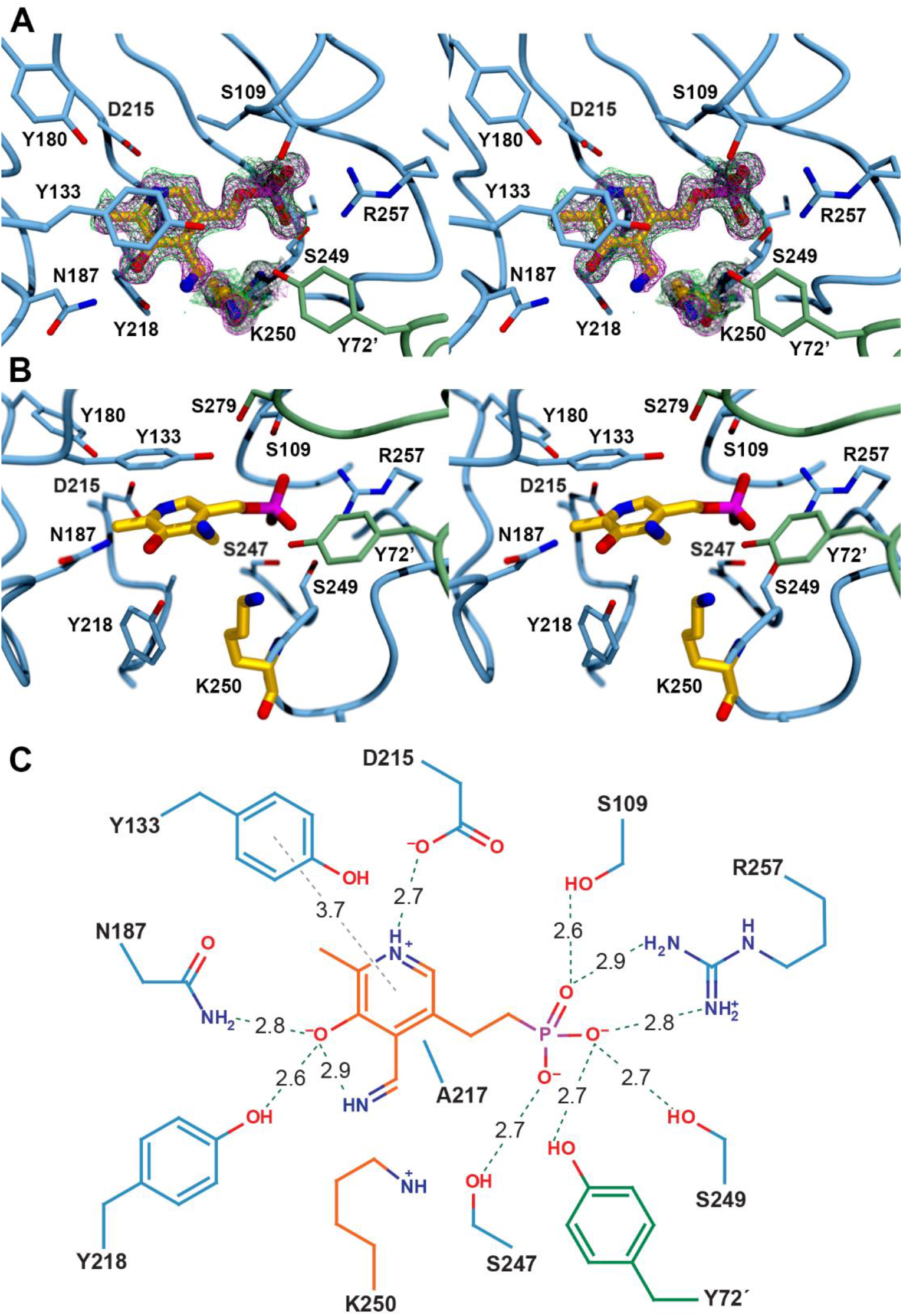
The 2|F_o_|-|F_c_| (magenta mesh) and simulated annealing composite omit (green mesh) electron density maps are shown for the PMP at 1.2 σ (map radius 2.5 Å) in a stereoview of the ShMppQ active site (A). A second stereoview of the active site (B) is shown without electron density and in a different orientation (rotated ~60 ° in the horizontal) to give a clearer picture of the PMP and catalytic lysine. A schematic representation of the active site (C) shows the interactions of active site functional groups with the cofactor and each other (associated distances are given in Å).

### Preliminary analysis of MppQ catalytic activity

The typical transamination reaction begins with the attack of the deprotonated amine of an amino acid on the C4’ imine of PLP to form the external aldimine. Hydrolysis of this adduct releases the keto-acid and leaves PMP in the enzyme active site. A different α-keto acid, often α-ketoglutarate, binds and reacts with PMP, forming a second external aldimine. Given the structural evidence that ShMppQ is an aminotransferase, and that it operates in the biosynthesis of L-End from L-Arg, we used steady state kinetics to assess the reaction of the enzyme with a number of amino group donor and acceptor.

In order to observe the aminotransferase activity of MppP, a coupled assay with lactate dehydrogenase (LDH) was designed. This assay relied on L-alanine as an amino group donor being converted to pyruvate, which is then reduced to lactate, while oxidizing NADH in the process (Scheme 3). Through following the decay of NADH at 340 nm, preliminary steady state kinetics data was collected. It was shown that MppQ catalyzes transamination of **7** and L-alanine (Figure 4), as well as its proposed physiological substrate, **6**, and L-alanine. The low K_M_ values indicates that MppQ has a high affinity for **7**. It is plausible that this serves a physiological role, as it would allow MppQ to recycle the “shunt product” **7**, back to L-arginine, thus providing MppP with more substrate and increasing the yield of L-End produced. The reactions of MppQ with 2KE and L-Ala manually quenched and analyzed using MS, have confirmed the substrate turnover (Figure 5, Figure 6).

**Scheme 3.**
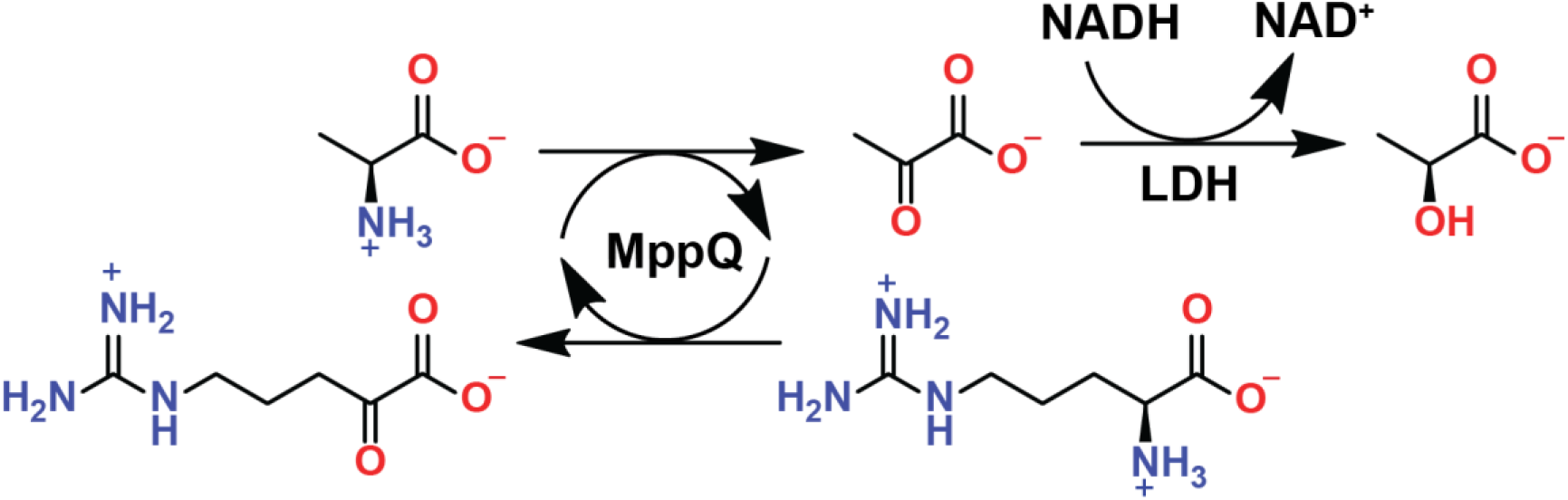
An assay of MppQ reaction with 7 and L-Ala, coupled with LDH reaction with pyruvate produced by MppQ.

**Figure 4.**
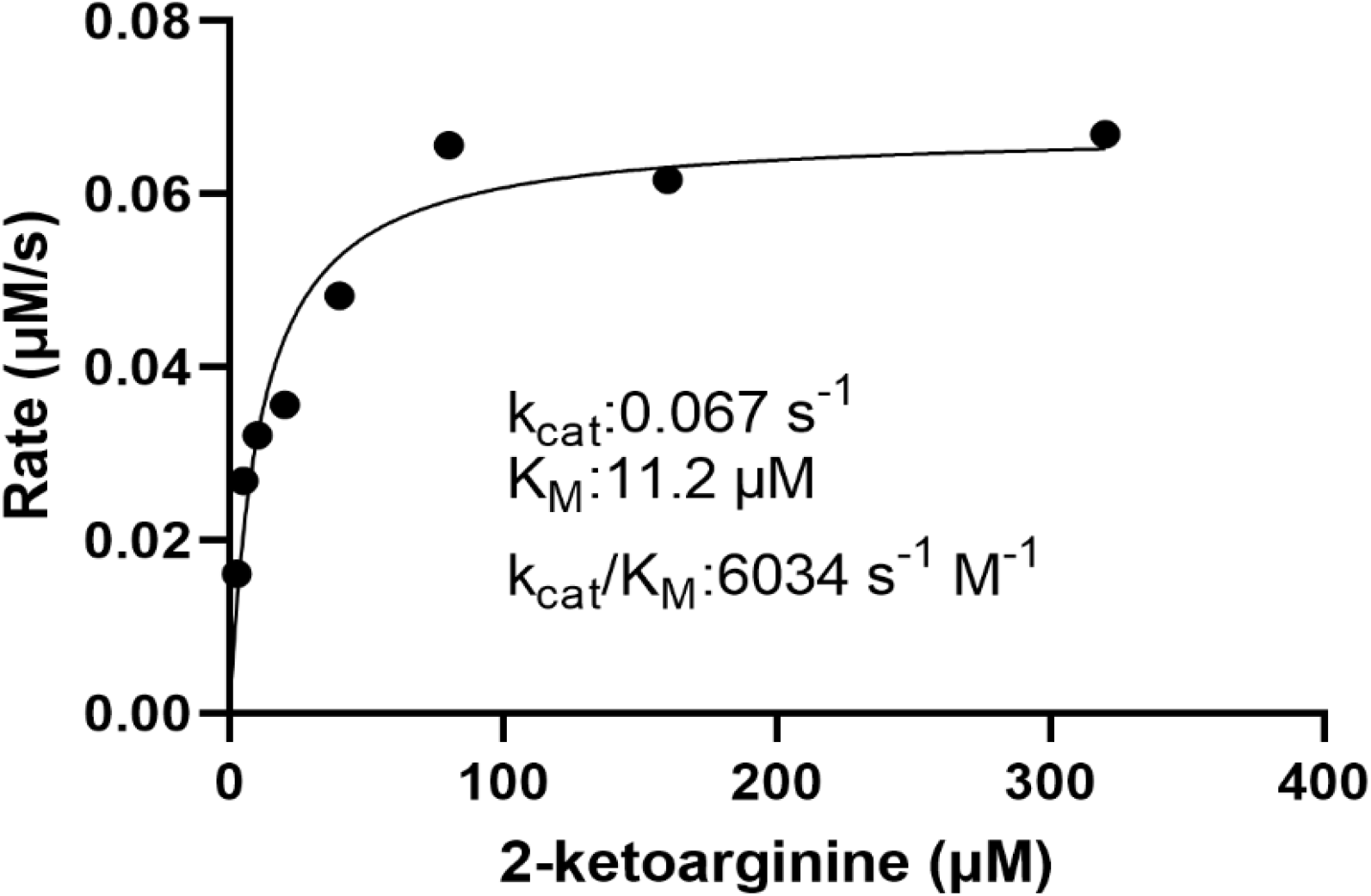
Michaelis-Menten curve for the reaction of ShMppQ with **7** and L-Ala.

**Figure 5.**
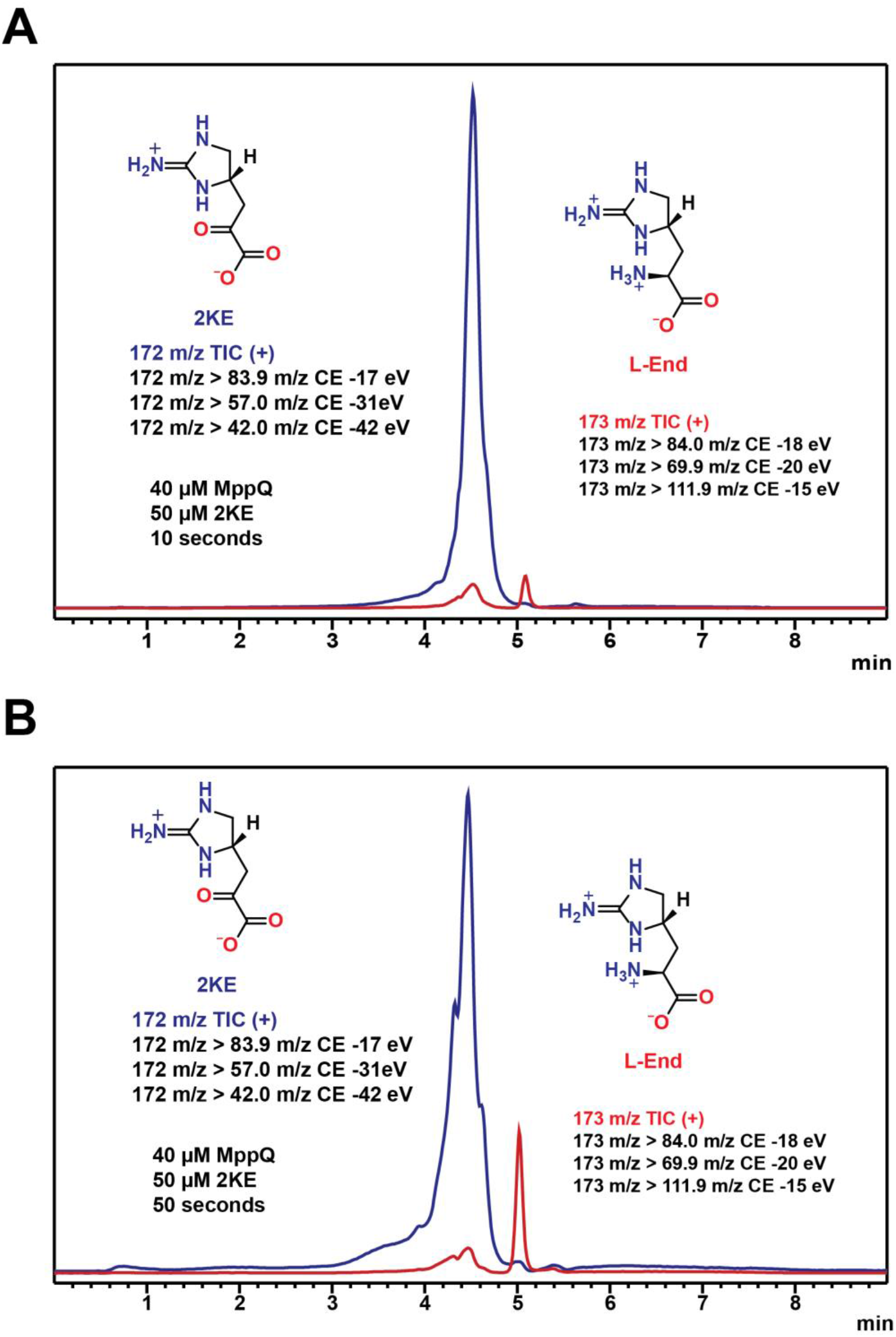
MS chromatograms of MppQ reaction with **5** and L-Ala at 10 (A) and 50 seconds (B).

**Figure 6.**
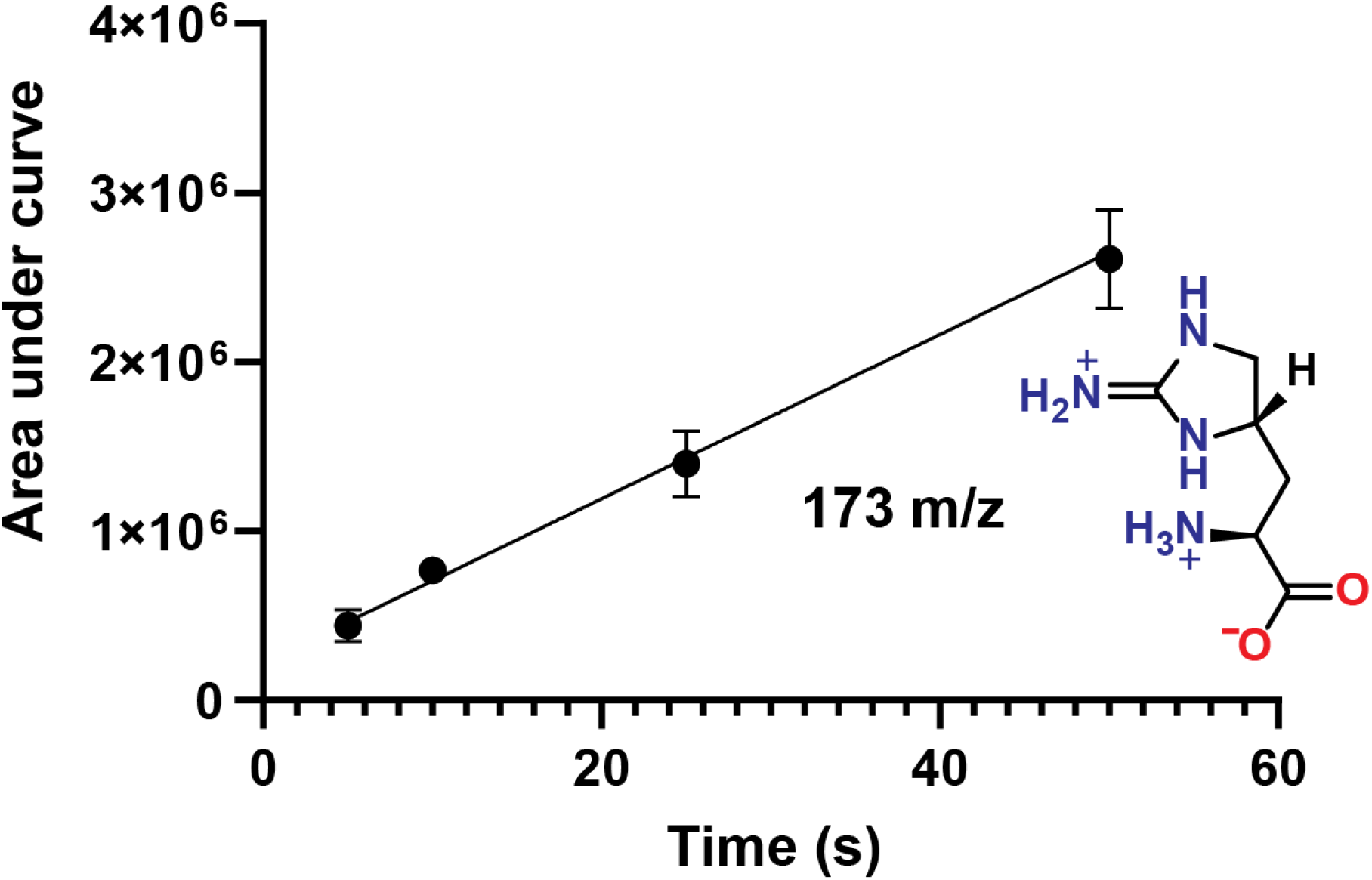
Production of 1 in a reaction of ShMppQ with 6 and L-Ala. The peaks corresponding to all fragments generated from 173 m/z precursor ion were integrated and areas under the curve plotted on the y-axis.

However, L-Ala is a very poor substrate (K_M_ > 50 mM, data not shown), leading to difficulty in obtaining accurate kinetic parameters. Previous HPLC-based end-point assays have indicated that MppQ reacts with L-Arg and glyoxylate, and that this reaction proceeds more rapidly than the analogous reaction with pyruvate as the amino group acceptor. However, it appears that this reaction is irreversible, as the reaction of ShMppQ with **7** and glycine produced no L-Arg (Figure 7).

**Figure 7.**
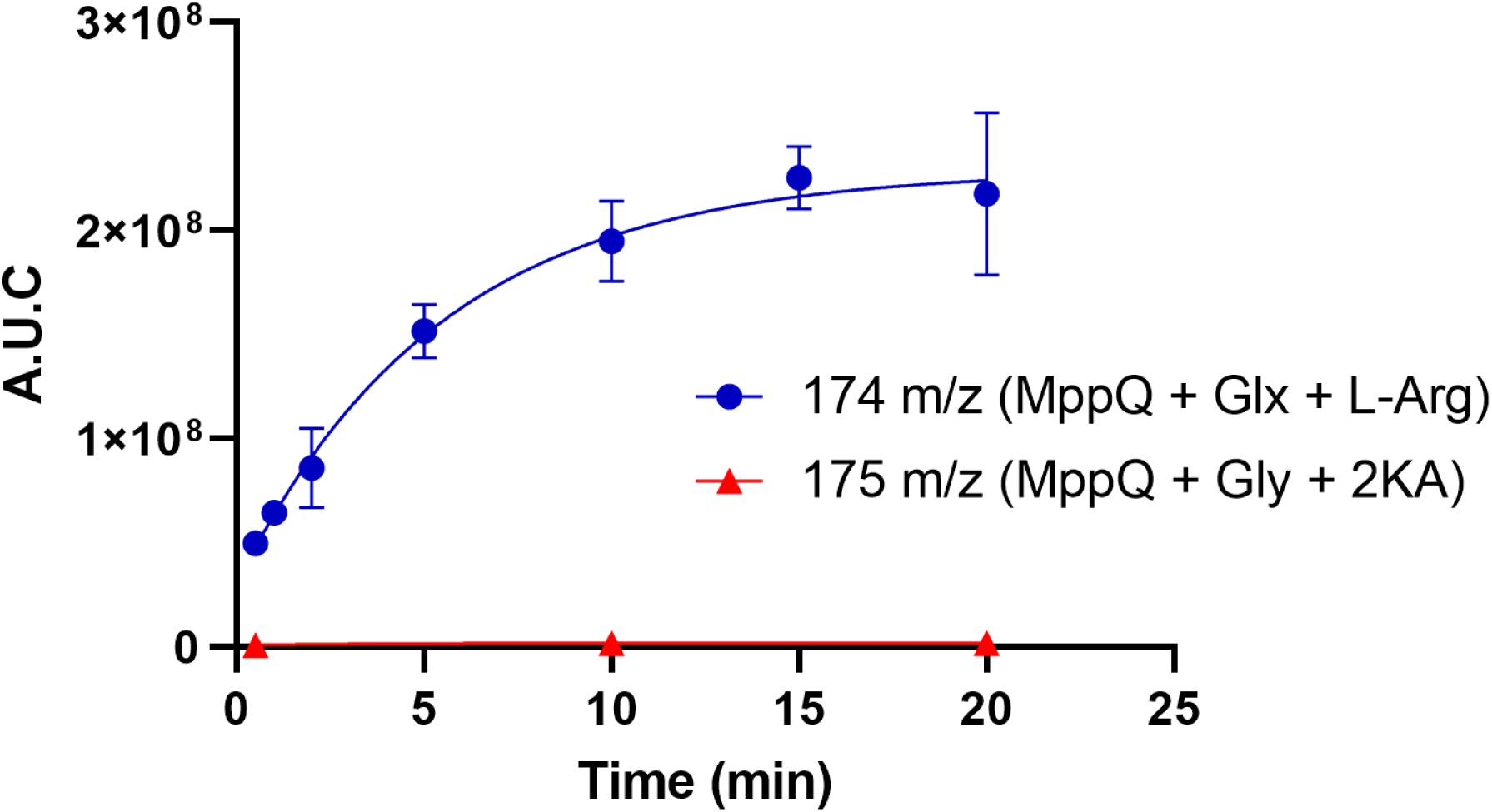
Production of 2KA (174 m/z) and lack of produced L-Arg (175 m/z) in reactions of MppQ with glyoxylate and glycine, respectively. The coupling partner in each reaction was **7**.

This is most likely due to glycine’s lack of a side chain which makes it more difficult to be properly oriented for catalysis. Using MS-based end-point assays, different amino group donors were tested, and the best donor substrate was shown to be L-ornithine (Figure 8).

**Figure 8.**
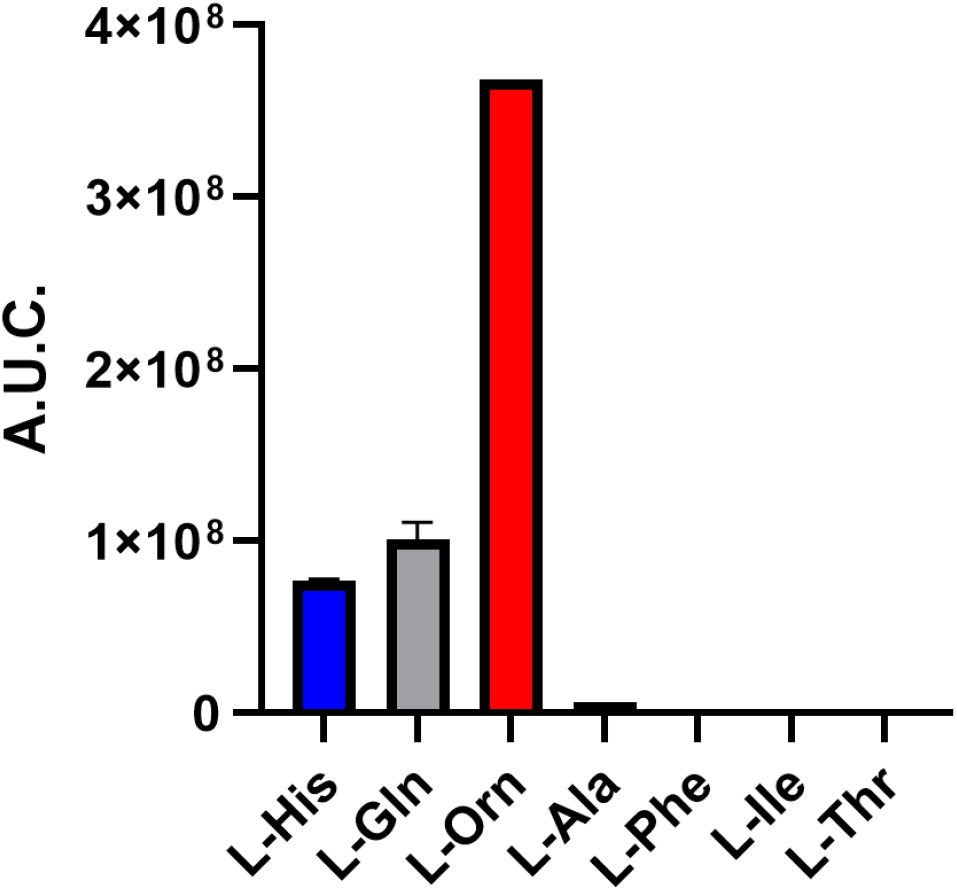
Reactivity of MppQ and **7** with various amino donor substrates.

The fact that the sidechain of L-ornithine possesses a charged amine may explain its exceptional activity with MppQ as an amino group donor since MppQ’s physiological substrate, **6**, possesses a positively charged iminoimidazolidine moiety.

