## Supplemental Information for "Structural and Preliminary Biochemical Characterization of MppQ, a PLP-Dependent Aminotransferase from *Streptomyces hygroscopicus*"

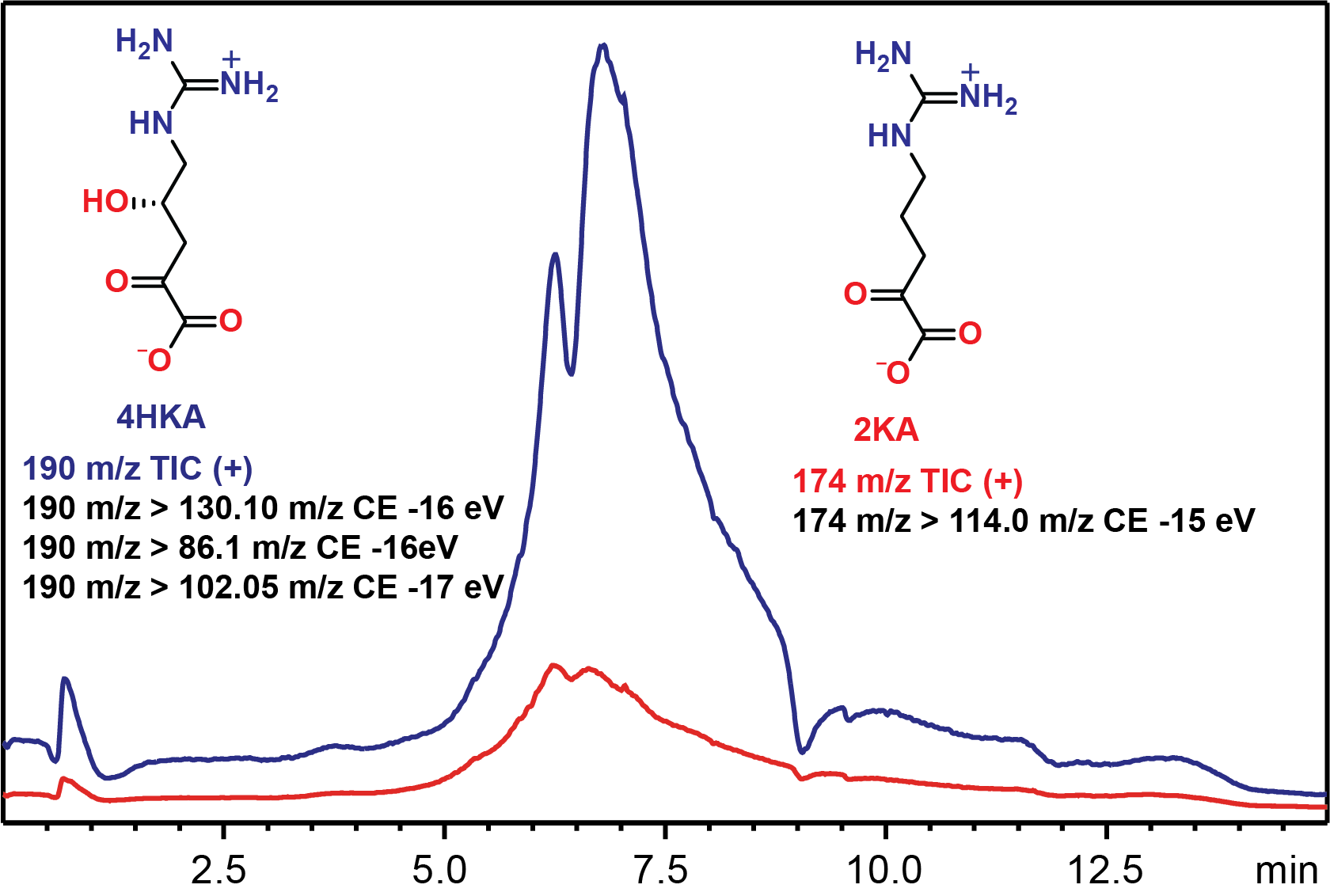

**Figure** **S1.** Chromatogram of enzymatically produced mixture of 4(S)HKA and 2KA in 20 mM ammonium acetate, pH 7.0.

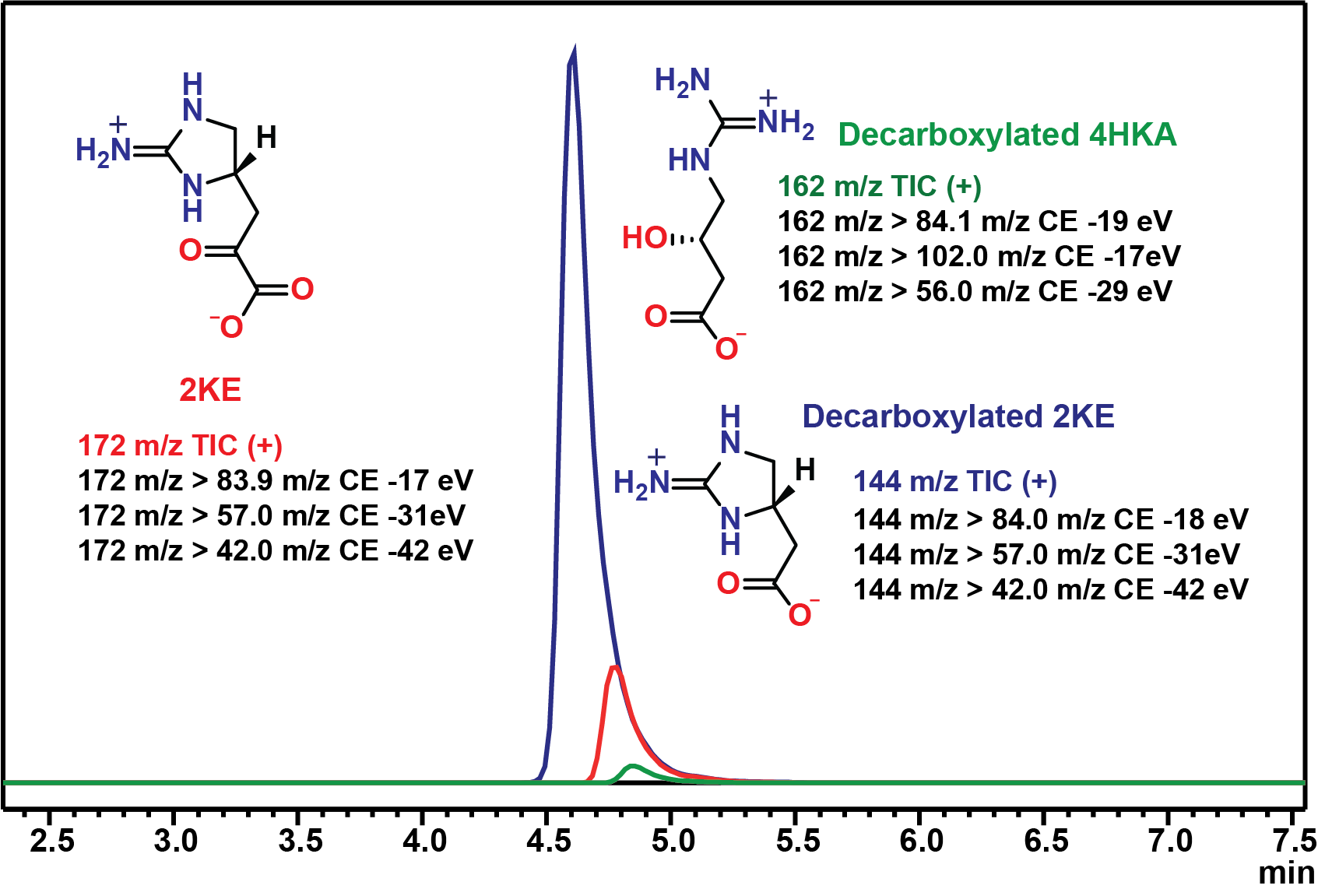

**Figure S2.** Chromatogram of enzymatically produced 2KE along with reaction by-products. The majority of 2KE appears to have been decarboxylated during the drying process, significantly decreasing product yield.
